# Hormonal and neural correlates of care in active versus observing poison frog parents

**DOI:** 10.1101/765503

**Authors:** Eva K. Fischer, Lauren A. O’Connell

## Abstract

The occasional reversal of sex-typical behavior suggests that many of the neural circuits underlying behavior are conserved between males and females and can be activated in response to the appropriate social condition or stimulus. Most poison frog species (Family Dendrobatidae) exhibit male uniparental care, but flexible compensation has been observed in some species, where females will take over parental care duties when males disappear. We investigated hormonal and neural correlates of sex-typical and sex-reversed parental care in a typically male uniparental species, the Dyeing Poison Frog (*Dendrobates tinctorius*). We first characterized hormone levels and whole brain gene expression across parental care stages during sex-typical care. Surprisingly, hormonal changes and brain gene expression differences associated with active parental behavior in males were mirrored in their non-caregiving female partners. To further explore the disconnect between neuroendocrine patterns and behavior, we characterized hormone levels and neural activity patterns in females performing sex-reversed parental care. In contrast to hormone and gene expression patterns, we found that patterns of neural activity were linked to the active performance of parental behavior, with sex-reversed tadpole transporting females exhibiting neural activity patterns more similar to those of transporting males than non-caregiving females. We suggest that parallels in hormones and brain gene expression in active and observing parents are related to females’ ability to flexibly take over parental care in the absence of their male partners.

Graphical abstract

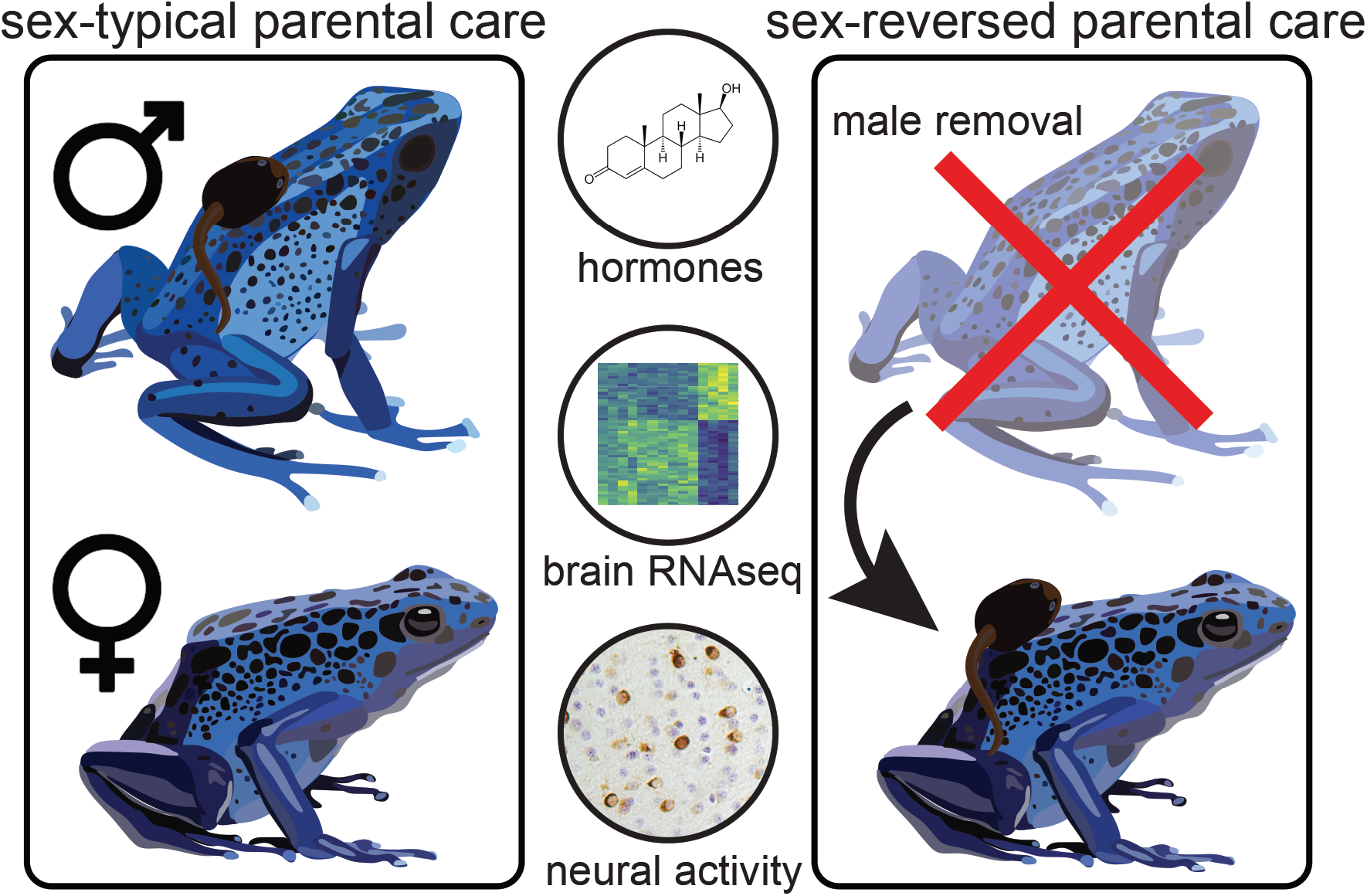

## Introduction

Sex differences in behavior and neuroendocrine function are a fundamental feature of most animals. Although sex-reversed behavior is relatively uncommon in adult animals, its existence suggests that neural circuits underlying behavior are conserved across sexes and can be activated under some circumstances, despite physiological and morphological differences established during development (Pereira and Ferreira, 2016). Exploring sex-reversed behavioral flexibility provides potential inroads to understanding how sex-specific behavioral patterns are coded in the brain, the conditions under which these patterns can be altered or reversed, and how underlying mechanisms are environmentally, developmentally, and evolutionarily tuned to give rise to sex-specific behavior.

Parental care is a behavior for which many species exhibit sexually dimorphic patterns. Parenting is ubiquitous across the animal kingdom, having arisen independently multiple times and taking different forms across species (Clutton-Brock, 1991; Royle et al., 2012). Species vary in the extent to which males and/or females provide care to offspring, and care behavior may be similar between sexes or sex-specific. Aside from sex-typical parental behaviors, some biparental species show flexibility in behavior, where a parent will compensate care effort when their partner disappears. This behavioral flexibility has been described in beetles (Smiseth et al., 2005), fish (Lavery and Reebs, 2010; Smiseth et al., 2005), birds (Harrison et al., 2009), and primates (Harrison et al., 2009; Zahed et al., 2009). However, less is known about flexibility in uniparental species where the normally non-parenting sex will occasionally care for offspring. Moreover, despite the phylogenetic diversity in parental behavior, mechanistic neuroendocrine research is heavily biased towards rodents that are female uniparental in the wild, although some can exhibit biparental care under laboratory conditions (Kohl et al., 2017; Numan and Insel, 2006). Maternal care evolved at the base of the mammalian lineage and shows little flexibility given the dependence of offspring survival on lactation. As a result, studies of male parental care come almost exclusively from biparental systems (Rilling and Mascaro, 2017), where parental behavior cannot easily be dissociated from pair bonding.

A fundamental question left open by heavily female-biased parental care research is how sex differences in offspring care arise and evolve. Despite marked sex differences in behavior, the neural circuits governing parental care appear largely conserved across sexes (Kohl and Dulac, 2018; Pereira and Ferreira, 2016), as are the expression patterns of key neuromodulators, such as galanin (Kohl et al., 2018; Wu et al., 2014) and prolactin (Angelier et al., 2016; Angelier and Chastel, 2009; Hashemian et al., 2016). Some sex-typical care behaviors are linked to differences in circulating hormones (Adkins-Regan, 2005), expression of key signaling molecules (Albers, 2015), and variations in receptor distributions in the brain (Dumais and Veenema, 2016). Nonetheless, the existence of flexible parenting suggests that underlying neural circuits are largely conserved and that there are alternative routes of access to their activation (Kohl and Dulac, 2018; Pereira and Ferreira, 2016). It remains an open question how and in response to which cues the modulation of these circuits leads to sex-reversed behavior.

Dendrobatid poison frogs show diversity in parental care across closely related species, including male uniparental, female uniparental, and biparental care (Summers and Tumulty, 2013). Parental care in poison frogs involves defense, hydration, and cleaning of embryos during development, and transportation of tadpoles piggyback to pools of water upon hatching (Brown et al., 2010; Pröhl and Hödl, 1999; Weygoldt, 1987). Importantly, males and females exhibit similar care behaviors within and across species, and both male and female care occur with and without pair bonding in this clade (Summers and Tumulty, 2013). Moreover, a number of species exhibit plasticity in parental care behavior, where the typically non-parenting sex will occasionally perform parental duties. For example, in brilliant-thighed poison frogs (*Allobates femoralis*) males typically perform tadpole transport, but females will do so if their mate disappears in both the wild and the laboratory (Ringler et al., 2015). Conversely, in the strawberry poison frog (*Oophaga pumilio*), females typically perform tadpole transport, but males have been observed doing so as well (Killius and Dugas, 2014). In poison frogs, the diversity of behavioral care strategies *between* species combined with behavioral flexibility *within* species affords a unique opportunity to identify physiological, neural, and molecular contributions to parental care and its evolution (Roland and O’Connell, 2015). Moreover, male parental care is ancestral in the clade (Summer et al., 1999), providing an important evolutionary comparison for mammalian systems in which female care is ancestral and male uniparental care is absent.

In the current study, we took advantage of behavioral flexibility in the typically male uniparental poison frog *Dendrobates tinctorius* to characterize hormonal and neural correlates of sex-typical and sex-reversed parental care. We first characterized hormone levels and whole brain gene expression during sex-typical parental behavior across egg attendance and tadpole transport care stages. We then took advantage of behavioral plasticity in female *D. tinctorius* to compare hormonal changes and neural activity associated with parental care in males and females during sex-typical and sex-reversed care. Flexible parenting in poison frogs provides an exciting opportunity to explore how sex-specific behavioral patterns are coded in the brain and how neuroendocrine mechanisms are evolutionarily tuned to give rise to sex- and species-specific variations in parental care.

## Methods

### Animals husbandry

All *D. tinctorius* used in the study were sexually mature individuals housed in our breeding colony. Animals were housed in 18×18×18 inch glass terraria (Exoterra, Rolf C. Hagen USA, Mansfield, MA) with sphagnum moss, live philodendrons, coconut shelters with petri dishes for egg laying, and a single water pool for tadpole deposition. Frogs were kept on a 12:12 hour light cycle and fed wingless Drosophila fruit flies dusted with vitamin supplements three times weekly. All procedures were approved by the Harvard University Animal Care and Use Committee (protocol no. 12-10-1).

### Tissue collection

Animals were housed in breeding pairs to ensure parental/partner identity. To control for potential effects of parental experience, we allowed all pairs to successfully rear at least one clutch from egg-laying through tadpole transport prior to the study. For the ‘no care’ group, we collected frog pairs between parental bouts when they were not caring for eggs or tadpoles (N=8 pairs for hormones, N=5 pairs for RNAseq). For the ‘egg care’ group, we collected frogs one week after they laid a clutch of eggs (N=7-10 pairs for hormones, N=5 pairs for RNAseq). We collected only frogs that were caring for eggs, as evidenced by the healthy development of embryos. For the ‘tadpole transport’ group, we checked daily for transport behavior and collected tissue when we found frogs actively transporting (N=7-8 pairs for hormones, N=5 pairs for RNAseq). For all behavioral groups we simultaneously collected tissue from parental males and their non-caregiving female partners.

We induced sex-reversed tadpole transport in *D. tinctorius* females by removing males (the typically transporting sex) a few days before tadpoles hatched. Once hatched, tadpoles are typically transported within a few days and un-transported tadpoles do not survive more than a week due to desiccation (*personal observations*). We allowed females a maximum of seven days to transport their hatched tadpoles. When we found a transporting female, we collected tissue from that female (N=8) and a time-matched non-transporting control female from the same treatment (i.e. male removed with hatched tadpoles but who did not perform tadpole transport; N=7).

The duration of tadpole transport is variable, lasting hours in both the wild and the laboratory (Pašukonis et al., 2019; Rojas, 2014). We were unable to pin-point the initiation of transport behavior and collected samples at a range of transport time points (>30 minutes and <2 hours), an approach we have used successfully in *D. tinctorius* and other poison frog species (Fischer et al., 2019b). Given this experimental constraint, we interpret our findings as reflective of neuroendocrine differences associated with broadly defined parental care stages (i.e. egg care, tadpole transport) rather than the initiation of particular aspects of behaviors performed during these distinct parental stages. Given variation in the time of sample collection relative to the onset of transport behavior, our findings likely represent conservative estimates of hormone and gene expression differences associated with parental behavior, with the potential for additional and/or other differences mediating changes that occur at smaller time scales.

Tissue was collected in an identical manner for all behavioral groups. Frogs were rapidly captured, anesthetized with a 20% benzocaine gel, weighed and measured, and euthanized by rapid decapitation. This entire process took less than 5 min. We collected trunk blood into tubes containing heparin. We centrifuged blood for 5 min at 5000 x g at 4°C and aliquoted plasma into new microcentrifuge tubes that were stored at −80°C until hormone analysis. For RNA sequencing of male and female brains across parental care stages, whole brains were dissected, flash frozen in liquid nitrogen, and stored at −80°C until further processing. For immunohistochemical detection of neural activity, brains were placed into 4% paraformaldehyde at 4°C overnight and then transferred to a 30% sucrose solution for cryoprotection. Once dehydrated, brains were embedded in Tissue-Tek® O.C.T. Compound (Electron Microscopy Sciences, Hatfield, PA, USA), rapidly frozen, and stored at −80°C until cryosectioning.

### Hormone quantification and analysis

We assayed hormone levels for cortisol (ADI-900-071), testosterone (ADI-900-065), estradiol (ADI-900-008), and prostaglandin (ADI-900-069) using EIA immunoassay kits (Enzo Life Sciences, Farmingdale, NY). We re-suspended 7µl of plasma in 203µl of the appropriate assay buffer and ran each sample in duplicate as per manufacturer’s instructions.

Because of over-dispersion and non-normality in the data, we log-transformed all hormone measurements and used transformed data in statistical analyses. We used linear mixed models to examine the effects of sex, behavioral group (no care, egg care, tadpole transport), and their interaction on hormone levels. Body size (snout-vent length) was included as a covariate in all models. We ran separate models for each hormone and used Tukey corrected *post hoc* contrasts to examine pairwise differences between behavioral groups. We followed the same general analysis procedures for the sex-reversed transport dataset but included only a main effect of behavioral group (sex-reversed transport vs no transport control) as all focal individuals were female. Mixed models were run in SAS (SAS 9.4; SAS Institute for Advanced Analytics) and data visualization was done using the ggplot2 package (Wickham, 2009) in R (version 3.5.0; the R Foundation for Statistical Computing).

### Library construction and sequencing

Flash frozen brains were placed in Trizol (Life Technologies, Grand Island, NY) and RNA was extracted according to manufacturer instructions. RNA quality was confirmed on an Agilent 2100 Bioanalyzer (Santa Clara, CA, USA) where all samples had a RIN (RNA Integrity Number) score >7; one sample (a male transporter) had a RIN score below 5 and was excluded from the further tissue processing and gene expression analysis. Poly-adenylated RNA was isolated from each sample using the NEXTflex PolyA Bead kit (Bioo Scientific, Austin, TX, USA) following manufacturer guidelines. Lack of contaminating ribosomal RNA was confirmed using an Agilent 2100 Bioanalyzer. Strand specific libraries for each sample were prepared using the dUTP NEXTflex RNAseq kit (Bioo Scientific), which includes a magnetic bead-based size selection. Libraries were pooled in equimolar amounts after library quantification using both quantitative PCR with the KAPA Library Quantification Kit (KAPA Biosystems, Wilmington, MA, USA) and the fluorometric Qubit dsDNA high sensitivity assay kit (Life Technologies), both according to manufacturer instructions. Libraries were sequenced on an Illumina HiSeq 2500 at the Bauer Sequencing Core at Harvard University.

### Transcriptome construction and annotation

We constructed *de novo* transcriptomes using a pipeline established in our laboratory (Caty et al., 2019). We amended sequencing errors using Rcorrector (Song and Florea, 2015) and trimmed raw reads using Trim Galore! (Babraham Bioinformatics, Babraham Institute) to remove Illumina adapters and restrict all reads to high-quality sequence. Following developer recommendations, we used a quality score of 33, a stringency of 5, and a minimum read length of 36 bp. Corrected, trimmed reads from all individuals were pooled prior to *de novo* transcriptome construction with Trinity (Grabherr et al, 2011; Haas et al, 2013). Our initial assembly contained 2,655,485 isoforms representing 2,272,431 presumptive genes.

We filtered the raw assembly using several approaches. We clustered overlapping contigs using CD-HIT-EST (http://weizhongli-lab.org/cd-hit/) and removed contigs that were smaller than 250 bp following clustering, leaving 1,479,791 total contigs. Preliminary annotations of this and other assemblies from closely related poison frog species revealed a high percentage of contaminant (i.e. non-vertebrate) sequences. To remove non-vertebrate (i.e. bacterial, fungal and other pathogen/parasite) contaminants, we annotated sequences using blastx queries with default parameters and an e-value cutoff of 10^−10^ against the SwissProt database (www.uniprot.org) and retained only those contigs with annotations to known vertebrate genes. Our final assembly contained 68,399 contigs representing 35,058 genes. Additional assembly annotation was done using Trinotate (Bryant et al, 2017) and assembly completeness was assessed using BUSCO (Simão et al, 2015). Final assembly completeness was estimated at 87% (37% duplicated; 3.5% fragmented, 9.0% missing). All high-powered computing for transcriptome assembly and filtering was performed on the Odyssey computing cluster supported by the FAS Science Division Research Computing Group at Harvard University. Our final, annotated *D. tinctorius* transcriptome assembly is available on DataDryad (https://doi.org/10.5061/dryad.3n5tb2rd0). Raw reads are available from the NCBI SRA (PRJNA601475).

### Read quantification and differential expression analysis

We mapped reads back to our final, filtered assembly and estimated their abundance using Kallisto (Bray et al, 2016) with default parameters. On average 54% of sequences per individual mapped back to our vertebrate only reference transcriptome. Read counts from all individuals were combined into a single matrix. We performed all subsequent analyses at the gene rather than transcript level as this has been demonstrated to give more reliable, interpretable results (Freedman et al., 2019).

We used principal components analysis (PCA) to characterize expression variance across all transcripts. We transformed raw transcript expression values with the variance stabilizing transformation in DESeq2 in R (Love et al., 2014) and performed PCA using the prcomp function in the R base package (version 3.5.0; The R Foundation for Statistical Computing). We tested for sex and behavioral group differences in principal components using the aov function in the R base package (version 3.5.0; The R Foundation for Statistical Computing).

We also used DESeq2 to perform differential expression analysis. We compared transcript expression among behavioral groups (no care, egg care, and tadpole transport) in a pair-wise fashion separately for males and females. Exploratory analyses found no significant group*sex interactions, and we therefore did not consider these effects further. We corrected p-values for multiple hypothesis testing and considered those transcripts with false discovery rate (FDR) corrected p-values <0.05 significantly differentially expressed. We examined overlap among differentially expressed transcripts and characterized overlapping genes as having expression differences in either the same or opposite directions based on log-fold expression differences.

We performed GO term enrichment analysis for set of genes differentially expressed during tadpole transport as compared to no care and/or egg care in males and females, as well as the set of differentially expressed genes overlapping between males and females, using ‘Biological Processes’ annotations in the topGO package in R (Alexa and Rahnenführer, 2009). Finally, we identified differentially expressed transcripts present in a collection of 158 transcripts of interest we previously compiled based on their known roles in parental care (Fischer et al., 2019b). Data visualizations were done using ggplot2 (Wickham, 2009) and pheatmap (www.rdocumentation.org/packages/pheatmap/versions/1.0.12) in R (version 3.5.0; The R Foundation for Statistical Computing).

### Neural activity differences between sex-reversed and control females

We performed tissue processing and analysis of neural activity in sex-reversed transporting and non-transporting control females in a manner identical to that previously described for three closely related species of poison frog, including *D. tinctorius* (Fischer et al., 2019b). Briefly, we sectioned brains into four coronal series at 14µm. To assess the level of neural activity across brain regions, we used an antibody for phosphorylated ribosomes (pS6; phospho-S6 Ser235/236; Cell Signaling, Danvers, MA, USA) and followed procedures standard in our lab for immunohistochemical staining with 3’,3’-diaminobenzadine (DAB). Ribosomes become phosphorylated when cells are active, and pS6 staining therefore represents a measure of ongoing neural activity (Knight et al., 2012). Detailed methodological descriptions are in (Fischer et al., 2019b).

Stained brain sections were photographed on a Leica DMRE connected to a QImaging Retiga 2000R camera at 20X magnification. We quantified labeled cells from photographs using FIJI image analysis software (Schindelin et al., 2012). Brain regions were identified using our dendrobatid brain atlas (Fischer et al., 2019b). We measured the area of candidate brain regions and counted all labeled cells in a single hemisphere for each brain region across multiple tissue sections. We quantified cell number in twelve brain regions known to mediate social decision-making across vertebrates (O’Connell and Hofmann, 2011): the nucleus accumbens, the basolateral nucleus of the stria terminalis, the habenula, the lateral septum, the magnocellular preoptic area, the medial pallium (homolog of the mammalian hippocampus), the anterior preoptic area, the suprachiasmatic nucleus, the striatum, the posterior tuberculum (homolog of the mammalian midbrain dopamine cells representing the ventral tegmental area and substantia nigra), the ventral hypothalamus, and the ventral pallidum.

We analyzed the relationship between parental behavior and pS6 neural activity to identify brain regions whose activity differed in sex-reversed transporting and control females. To test for differences in neural activation across brain regions, we used generalized linear mixed models with a negative binomial distribution appropriate for count data with unequal variances. We included behavioral group (sex-reversed tadpole transport vs non-caregiving control), brain region, and their interactions as main effects predicting the number of pS6-positive cells. Individual was included as a random effect. Brain region area was included as a covariate to control for body size differences between frogs, known size differences between brain regions, and rostral to caudal size/shape variation within brain regions. We explored main effects of group, sex, and regional differences in further detail using *post hoc* comparisons with Tukey adjustment for multiple hypothesis testing. We qualitatively compared neural activity differences during sex-reversed and sex-typical transport but did combine these data into a single analysis as sex-typical data are from a previously published study (Fischer et al., 2019b) and were collected and processed at a different time. We used model-estimated cell counts for data visualizations as these predicted cell numbers reflect values that have been adjusted for individual variation in body size, activity levels, and sampling. Statistical analyses were performed in SAS (SAS 9.4; SAS Institute for Advanced Analytics) and data visualizations in R (version 3.5.0; the R Foundation for Statistical Computing).

## Results

### Hormone differences across parental care stages and sexes

We quantified cortisol, testosterone, estradiol, and prostaglandin levels in parental male frogs and their non-caregiving female partners between parental bouts (no care), during egg care, and during tadpole transport (Fig. 1). Testosterone was overall higher in males (sex: F_1,41_=40.64, p<0.0001), but decreased in both sexes during tadpole transport (group: F_2,41_=10.39, p=0.0002). Cortisol differed marginally across care stages (group: F_2,40_=2.51, p=0.09), increasing in both males and females during tadpole transport. Estradiol was higher in females (sex: F_1,39_=66.62, p<0.0001) and showed no differences across care stages. Prostaglandin levels did not differ based on either sex or care stage.

**Figure 1.**
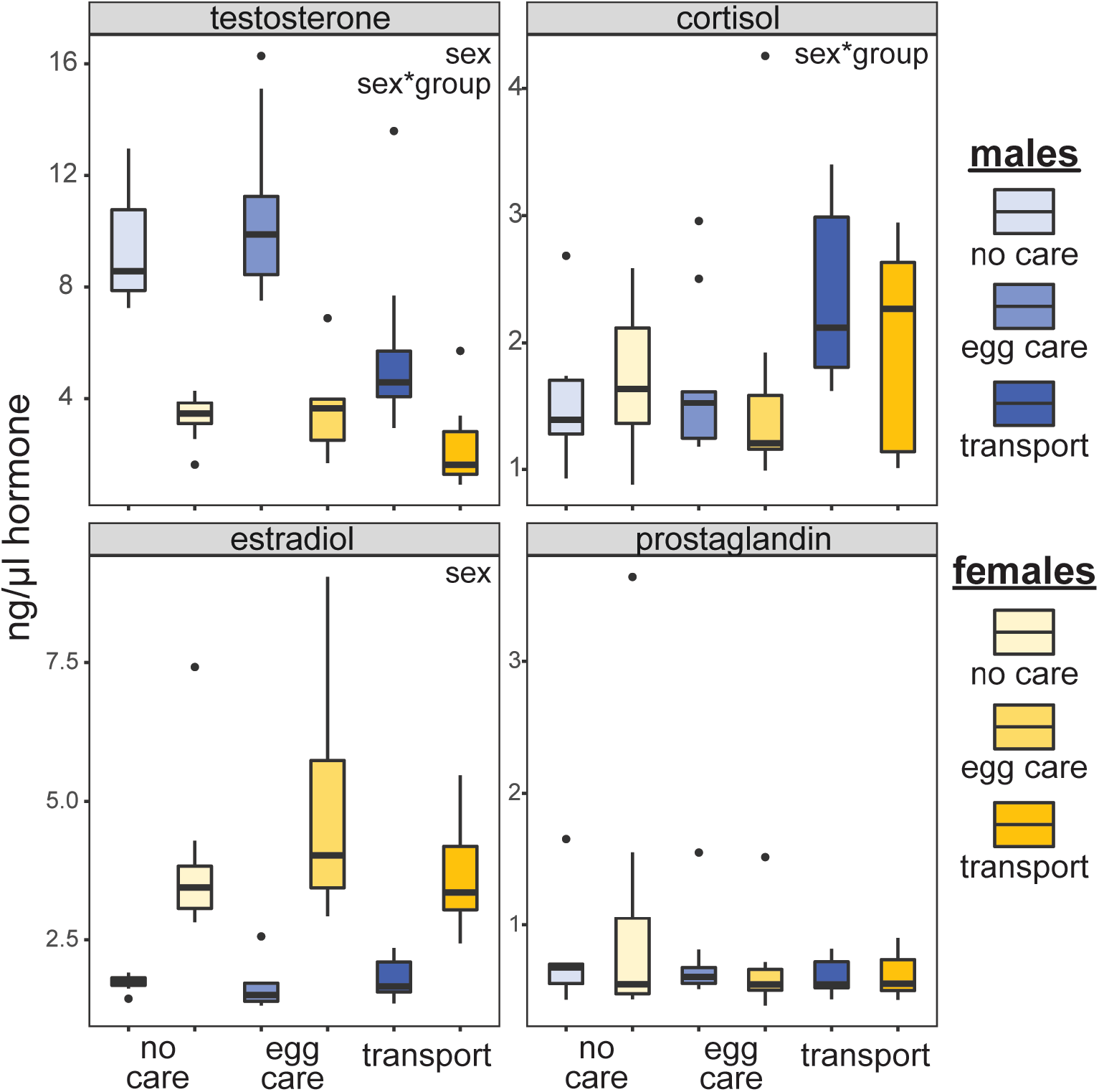
Hormone differences across care stages and sexes. Data are shown for males (blue) and their non-caregiving female partners (yellow) across care stages. Testosterone was overall higher in males but decreased in both transporting males and their non-caregiving female partners during tadpole transport. Conversely, cortisol increased in both sexes during tadpole transport. Estradiol was overall higher in females but showed no differences across care stages in either sex. Prostaglandin did not differ based on either sex or care stage. Significant main effects for each hormone are indicated in the top right corner of each plot.

### Brain gene expression patterns across parental care stages and sexes

We used principal component analysis to visualize overall brain gene expression variation between parental care stages and sexes. Principal component (PC) 1 explained 16%, PC2 11%, and PC3 8% of the overall variance in transcript expression. Tadpole transport samples separated significantly from no care and egg care samples along PC1 (F_2,23_=5.03, p=0.015) and PC3 (F_2,23_=6.79, p=0.004) (Fig. 2), but not PC2. We observed no significant separation of sex or group*sex along any of the first three PCs.

**Figure 2.**
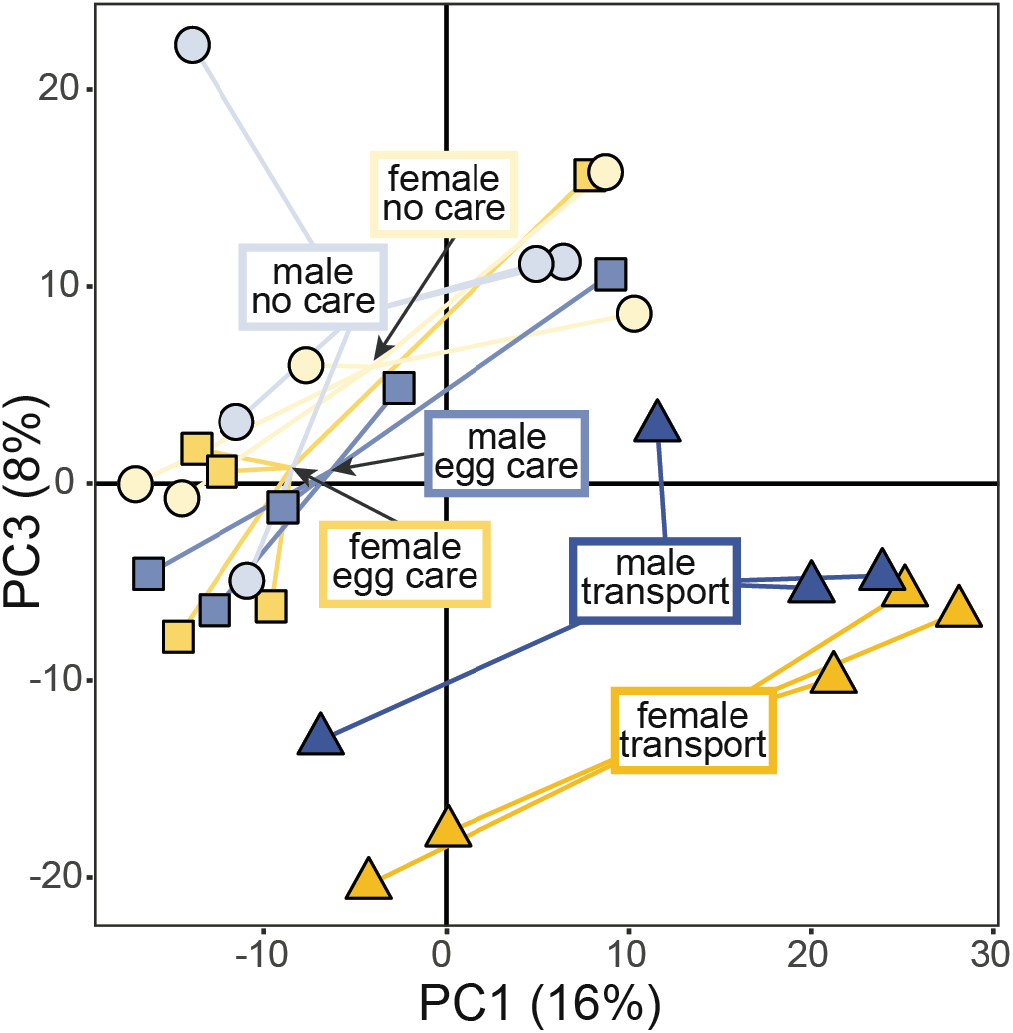
Principal component analysis of overall brain gene expression differences across care stages and sexes. Tadpole transport samples separate significantly from no care and egg care samples along PC1 and PC3 (darker colored circles cluster toward bottom right), but samples do not separate based on sex (no clustering of blue versus yellow circles). We found no sex and/or group differences in PC2 (11%). Percentages indicate the amount of expression variance explained by each PC.

Similar to patterns in overall brain gene expression, expression patterns among differentially expressed transcripts were most pronounced during tadpole transport as compared to either no care or egg care (Fig. 3). In males, only two transcripts were differentially expressed between no care and egg care, while 567 transcripts were differentially expressed between no care and tadpole transport, and 116 transcripts were differentially expressed between egg care and tadpole transport (Fig. 3A; Table S1). In females, only one transcript was differentially expressed between no care and egg care, while 1,114 transcripts were differentially expressed between no care and tadpole transport, and 1,492 transcripts were differentially expressed between egg care and tadpole transport (Fig. 3A; Table S2). Distinctive expression patterns associated with tadpole transport in both males and females are illustrated by sample clustering based on expression similarity (Fig. 2 & 3B).

**Figure 3.**
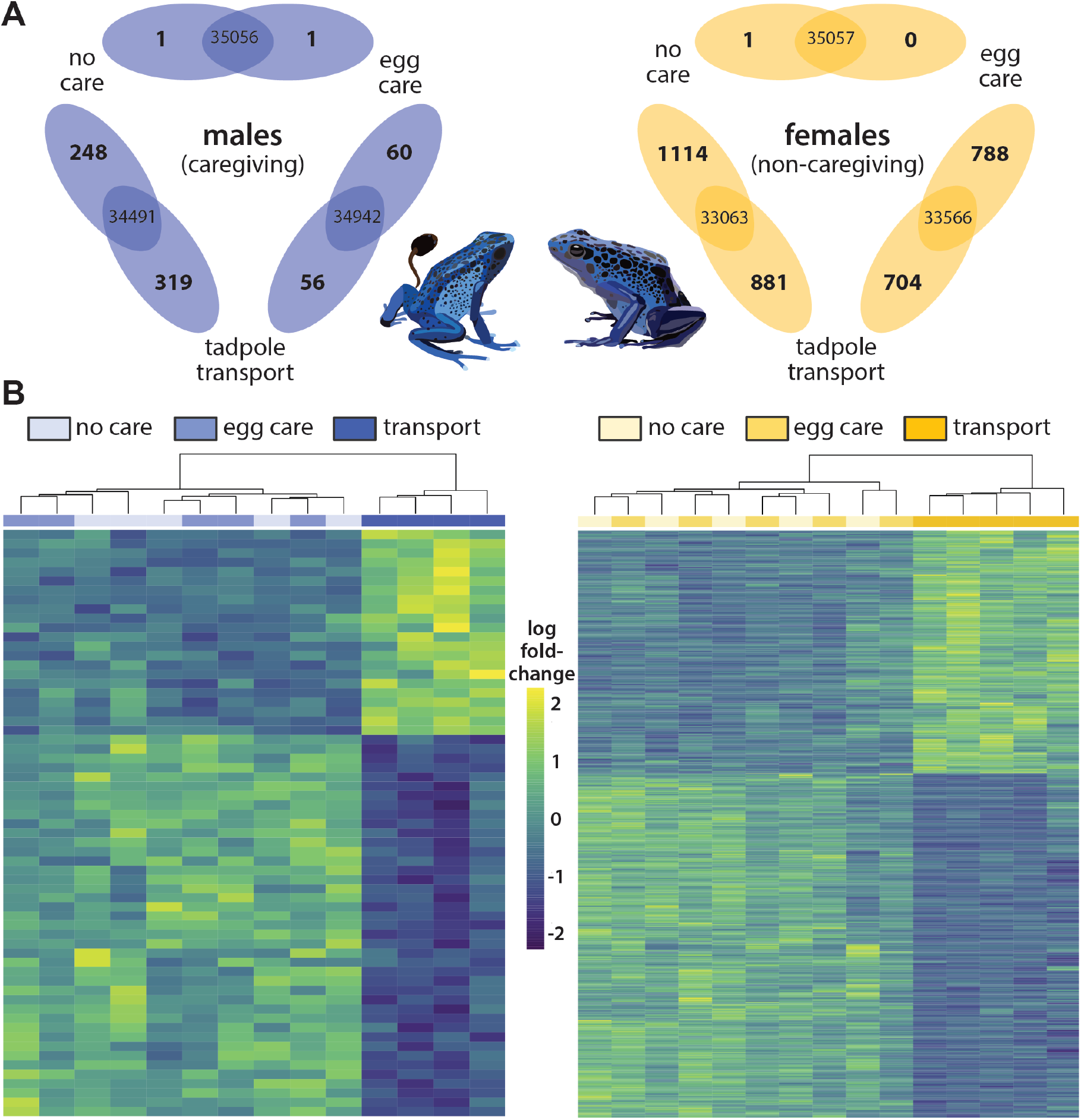
Differential brain gene expression across care stages and sexes. **(A)** Differential expression comparisons between care stages in males (blue) and females (yellow). Transcripts differentially expressed in pairwise comparisons between care stages are shown in Venn diagrams with the number of differentially expressed transcripts on the edges and the number of non-differentially expressed transcripts in the overlap. The number of genes closer to a given care stage indicates the number of transcripts that were significantly upregulated in that behavioral group. **(B)** Heat maps of differential transcript expression (rows) clustered based on expression similarity of individual frog samples (columns) across groups. Heat maps include those differentially expressed transcripts overlapping between pairwise comparisons of tadpole transport vs no care and tadpole transport vs egg care (N=63 transcripts in males, N=1,211 transcripts in females).

In males, 63 differentially expressed transcripts were overlapping between no care versus tadpole transport and egg care versus tadpole transport comparisons (Fig. 3B), 11% and 54% of the total number of differentially expressed transcripts in the respective comparisons. In females, 1,112 differentially expressed transcripts were overlapping between no care versus tadpole transport and egg care versus tadpole transport comparisons (Fig. 3B), 99% and 75% of the total number of differentially expressed transcripts in the respective comparisons. In both males and females, all overlapping transcripts showed expression differences in the same direction (i.e. either up or down regulated) in the tadpole transport group as compared to no care or egg care. Comparing these overlapping transcripts across sexes, 45 were shared between males and females (Table S3), again all with expression differences in the same direction.

We examined GO annotations for those genes differentially expressed during tadpole transport as compared to no care and/or egg care in males (N=63 transcripts) and females (N=1,112 transcripts), as well as in the set of differentially expressed transcripts overlapping between sexes (N=45 transcripts). Notable across all comparisons was an increase in metabolism related GO categories, including glycolytic processes and gluconeogenesis. Notable specifically in females was the enrichment of transcripts involved in translation and neurogenesis. Complete GO annotation information is in Supplemental Table S4.

Finally, we compared lists of all differentially expressed transcripts to a set of 158 transcripts of interest we previously compiled based on their known roles in parental care and/or their defining cell types in the hypothalamus (Fischer et al., 2019b). Of these previously identified transcripts, five were differentially expressed in males and 16 were differentially expressed in females (Table 1). Pro-neuropeptide Y and one cut domain family member 2 were differentially expressed during tadpole transport in both males and females.

**Table 1.** Differentially expressed transcripts with known roles in parental care. Relative expression difference during tadpole transport as compared to no care and egg care is indicated.

|  | transcript annotation | expression during tadpole transport |
| --- | --- | --- |
| **males** | bombesin | increased |
|  | cholecystokinin | increased |
|  | nitric oxide synthase, brain | decreased |
|  | one cut domain family member 2 | decreased |
|  | pro-neuropeptide Y | increased |
| **females** | calcitonin receptor | decreased |
|  | cerebellin-1 | Increased |
|  | chondroitin sulfate proteoglycan 5 | increased |
|  | galanin receptor type 2 | decreased |
|  | insulin-like growth factor 1 receptor | decreased |
|  | lysophosphatidic acid receptor 1-A | increased |
|  | myelin basic protein | increased |
|  | nitric oxide synthase, endothelial | increased |
|  | one cut domain family member 2 | decreased |
|  | POU domain, class 3, transcription factor 2-B | increased |
|  | pro-neuropeptide Y | increased |
|  | prolactin receptor | decreased |
|  | somatostatin-1 | increased |
|  | spermidine synthase | increased |
|  | synaptotagmin-4 | increased |
|  | TNF receptor-associated factor 4 | increased |

### Hormone levels and neural activity during sex-reversed transport in females

To further dissect sex-specific versus general neuroendocrine correlates of parental care, we characterized hormone levels and neural activity in *D. tinctorius* females exhibiting sex-reversed parental care following male mate removal (Fig. 4A). We found no differences in cortisol, testosterone, or estradiol levels between females performing sex-reversed tadpole transport and time-matched control females that also had their mates removed but did not transport their hatched tadpoles (Fig. 4B). In other words, both groups had hormone levels distinct from non-caregiving females during the no care and egg care stages, but similar to those of those of non-caregiving females with transporting male partners.

**Figure 4.**
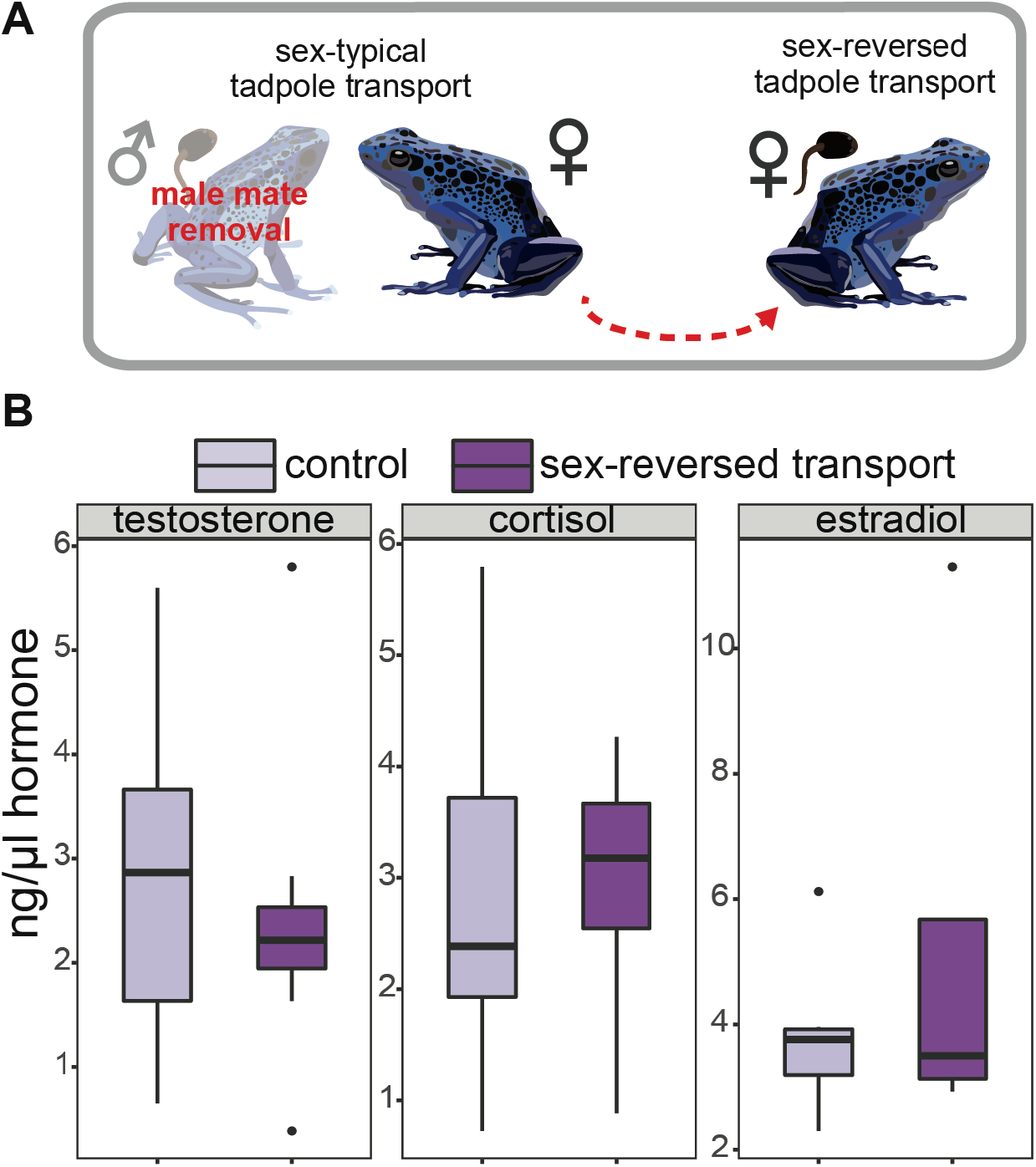
Hormone levels during sex-reversed tadpole transport. **(A)** We induced sex-reversed female tadpole transport via male mate removal. **(B)** We found no differences in testosterone, estradiol, or cortisol between females that performed sex-reversed tadpole transport following mate removal and time-matched control females that had their mates removed but did not transport their hatched tadpoles. Note that both groups had hormone levels similar to those on non-caregiving females with transporting male partners (see Fig. 1).

We next compared neural activity patterns in females performing sex-reversed tadpole transport and non-transporting control females. We found a significant interaction between behavioral group and brain region (group*region: F_1,1393_=5.10, p<0.0001; Table 2), and post hoc tests demonstrated differences in a number of brain regions (Table 3). Similar to patterns previously documented in tadpole transporting poison frogs independent of sex and species, sex-reversed transporting *D. tinctorius* females had increased neural induction in the preoptic area, and a marginally significant increase in medial pallium (Fig. 5A; Table 3). Additionally, increased neural activity in the anterior POA, magnocellular POA, habenula, and suprachiasmatic nucleus of sex-reversed *D. tinctorius* females was particularly notable as these regions were active in transporting male *D. tinctorius* but not their non-caregiving female partners (Fig. 5A & B).

**Table 2.** Summary of main statistical effects for neural activity differences between sex-reversed transporting females and non-transporting control females.

|  | df | F value | p value |
| --- | --- | --- | --- |
| behavioral group | 1,1393 | 7.34 | 0.0068 |
| region | 12,1393 | 58.43 | <0.0001 |
| behavioral group*region | 12,1393 | 5.10 | <0.0001 |

**Table 3.** Posthoc comparisons for behavioral group differences in neural activity by brain region.

| brain region | df | F value | p value |
| --- | --- | --- | --- |
| Acc | 1,1393 | 8.98 | **0.0028** |
| BST | 1,1393 | 0.28 | 0.5977 |
| DV/VP | 1,1393 | 1.99 | 0.1589 |
| H | 1,1393 | 7.49 | **0.0063** |
| Ls | 1,1393 | 0.03 | 0.8709 |
| Mgv | 1,1393 | 10.78 | **0.0011** |
| Mp | 1,1393 | 3.51 | 0.0611 |
| aPOA | 1,1393 | 15.91 | **<.0001** |
| mPOA | 1,1393 | 18.04 | **<.0001** |
| SC | 1,1393 | 4.61 | **0.0319** |
| Str | 1,1393 | 5.15 | **0.0234** |
| TP | 1,1393 | 0.00 | 0.9743 |
| VH | 1,1393 | 2.35 | 0.1254 |

**Figure 5.**
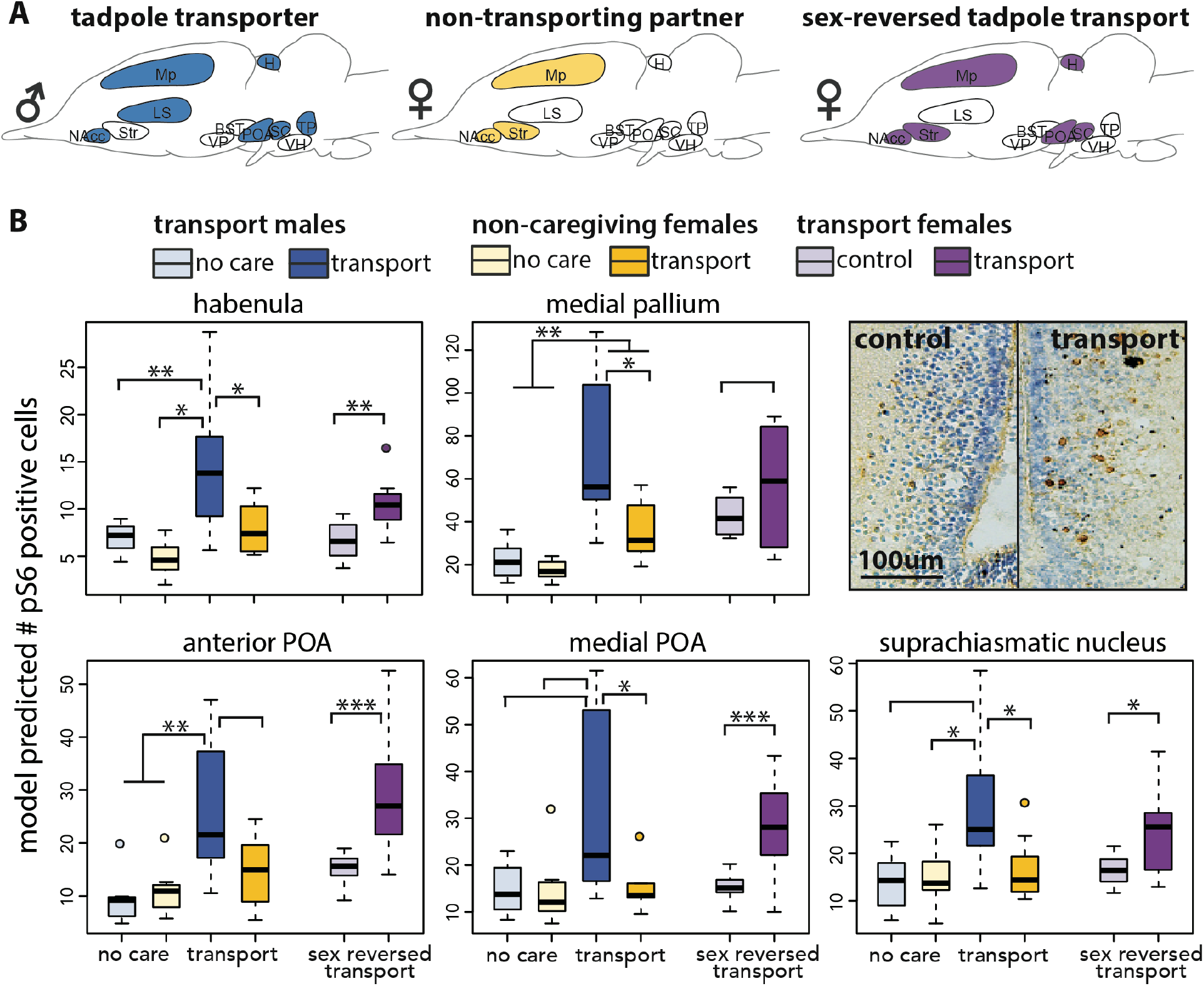
Neural activity during sex-reversed tadpole transport. **(A)** Schematics showing brain regions with significant neural activity differences during tadpole transport. Patterns of neural activity in transporting females (purple) were more similar to those in transporting males (blue) than their non-caregiving female partners (yellow). **(B)** Detailed results for brain regions associated with the active performance of tadpole transport in males and sex-reversed females. Representative micrographs of pS6 staining (brown) with cresyl violet nuclear staining (purple) from the aPOA are shown. P-values: no asterisk < 0.1, * <0.05, ** <0.01, *** <0.001. Abbreviations: Mp = medial pallium (homolog of the mammalian hippocampus), aPOA = anterior preoptic area, mPOA = magnocellular preoptic area. Sex typical data are from Fischer et al. (2019) and are shown for comparison but were not re-analyzed.

## Discussion

We compared hormone levels, whole brain gene expression, and neural activity across care stages in parental males, their non-caregiving female partners, and behaviorally sex-reversed females of the typically male uniparental poison frog *Dendrobates tinctorius*. During sex-typical care, hormone and gene expression signatures were most distinct during the tadpole transport stage of parental care in both males and females, even though females are not directly involved in sex-typical parental care. To explore this seeming disconnect between underlying neuroendocrine correlates and behavior, we took advantage of behavioral flexibility in this species to characterize hormone levels and neural activity of females performing sex-reversed tadpole transport. We suggest that parallelism in hormonal and gene expression patterns, but not neural activity, across males and females during sex-typical behavior is related to females’ monitoring of their partners’ behavior and ability to flexibly take over parental care in the absence of their male partners.

### Parallel hormonal shifts in active and observing parents

Hormones play conserved roles in physiology (e.g. Costantini et al., 2011; McCormick and Bradshaw, 2006; Wada, 2008) and behavior (e.g. Adkins-Regan, 2005; Brown, 1985; Young and Crews, 1995) across vertebrates, including in the induction and maintenance of parental behavior. Nonetheless, the precise nature of hormonal influences on parental care can differ by sex, species, and clade, with variations arising from behavioral and physiological differences linked to evolutionary history, life history, and patterns of care (reviewed in Adkins-Regan, 2005). While there are a few studies specifically examining the neuroendocrine basis of parental care in amphibians (see below), research on sexual and aggressive behavior suggests a conserved hormonal influences among frogs and other vertebrates (Heatwole and Carroll, 2000; Moore et al., 2005; Norris and Lopez, 2010; Wilczynski et al., 2005).

A hormonal shift associated with parental care that largely transcends sex, species, and life history differences is the increase in glucocorticoids (cortisol or corticosterone) observed during parental effort (da Silva Mota et al., 2006; Magee et al., 2006; Ouyang et al., 2011; Storey, 2000). Increases in cortisol/corticosterone may be related to the increased behavioral demands (e.g. offspring care and defense), and/or metabolic demands (e.g. increased foraging and offspring provisioning) associated with parental care in most species. As compared to non-parental frogs, we observed increased cortisol levels in males transporting tadpoles, but not males caring for eggs. Increased cortisol during the tadpole transport phase of parental care could be associated with the more active, costly nature of this behavior. In the wild, tadpole transporting *D. tinctorius* traverse large distances to deposit tadpoles across multiple water pools (Pašukonis et al., 2019; Rojas, 2014). In both the wild and the laboratory, transport of a single tadpole can last for hours and multiple tadpoles from the same clutch are typically transported individually, requiring repeated, lengthy trips outside the frog’s home range (Pašukonis et al., 2019). Increased cortisol during tadpole transport could therefore be associated with the increased energetic demands of tadpole transport, and/or behavioral adjustments required during transport (e.g. to mitigate trade-offs with other behaviors). Alternatively, changes in cortisol during tadpole transport but not egg care could arise if increased cortisol acts as a signal to initiate transport – for example triggered by tadpole hatching – rather than a response to performing transport. We cannot presently distinguish between facilitative versus responsive roles of cortisol and note that these functions are not mutually exclusive.

Changes in androgen levels are also associated with parental care across vertebrates (Gettler et al., 2011; O’Connell et al., 2012; Wingfield et al., 1990; Wynne-Edwards, 2001). Decreases in testosterone are extensively documented in parental males, and have also been observed in parental females (Fite et al., 2005; Fleming et al., 1997; Kuzawa et al., 2010). To our knowledge, the only previous study linking androgens to parental care in anurans found decreased testosterone in *Eleutherodactylus coqui* males during egg attendance and defense compared to non-parental individuals (Townsend and Moger, 1987). The general pattern of decreased androgen levels during parental effort may result from reduced testosterone facilitating parental care via a reduction in testosterone-mediated sexual and aggressive behavior (Adkins-Regan, 2005; Ketterson and Nolan, 1992). *D. tinctorius* males are not highly territorial or aggressive (Rojas and Pašukonis, 2019), and neither males nor females exhibit changes in territoriality or aggression associated with parental status, although this has not been tested experimentally. However, as non-seasonal breeders, short-term physiological shifts could be important in balancing reproductive and parental behaviors. As with cortisol, shifts in testosterone levels may be restricted to tadpole transport due to the more demanding, acute nature of this care stage as compared to the extended, less active nature of egg care. Alternatively, changes in testosterone may trigger tadpole transport by shifting behavior away from sexual pursuits. In brief, hormonal changes associated with parental care in *D. tinctorius* males are similar to those observed in other vertebrates, but additional research is necessary to disentangle causal versus responsive hormonal shifts associated with behavior.

Although females are not typically involved in tadpole transport, increased cortisol and decreased testosterone in male transporters was mirrored in their non-caregiving female partners. These changes in female physiology could be a cue (i.e. male behavior) or timing (i.e. time since last mating) dependent shift linked to reproductive cycling. For example, the hormonal state of one individual can influence the hormonal state of their dyad partner in pair bonding species (Edelstein et al., 2015; Lehrman, 1965; Storey et al., 2000). Although *D. tinctorius* are not pair-bonding and pairs do not associate beyond mating in the wild (Rojas and Pašukonis, 2019), laboratory housing in breeding pairs may increase interactions between partners such that changes in male behavior influence female physiology. Alternatively, parallel hormonal shifts across sexes may be related to flexible parenting, priming females to perform tadpole transport if it becomes necessary for offspring survival. Indeed, hormonal correlations between males and females in new and expectant human parents have been suggested to prime fathers for offspring care (Storey et al., 2000).

In support of the priming hypothesis, we found no hormone differences between sex-reversed transporting females and non-transporting control females who also had their mates removed but did not transport their hatched tadpoles. Both groups had hormone levels distinct from non-caregiving no care and egg care females but resembling those of non-caregiving females with transporting male partners. Thus, observing female parents showed hormonal changes mirroring those of males during tadpole transport, but no additional changes in response to male mate removal and/or active performance of tadpole transport (see also below). This suggests a priming function of hormonal changes rather than hormonal responses to active performance of tadpole transport and/or male disappearance. Additional testing and hormone manipulations in the laboratory and the field will be required to definitively distinguish between these possibilities.

### Parallel brain gene expression patterns in active and observing parents

Whole-brain gene expression patterns paralleled hormonal patterns in that tadpole transport was the most distinct care stage in both males and females. Tadpole transport is an acute phase of care, with increased physical and cognitive demands on parents who must transport their clutch to suitable water pools within a relatively narrow time window following hatching. For these reasons, hormonal and transcriptomic differences during tadpole transport may be more pronounced than during egg care, which is a longer, less active care phase. If hormonal and transcriptional changes reflect primarily short-term rather than long-term behavioral shifts, our experimental design is more likely to detect differences associated with tadpole transport (lasting hours) than those associated with egg care (lasting weeks). Some of the changes associated with tadpole transport could also represent time dependent or physiology dependent cues to initiate a new reproductive bout, rather than being associated directly with transport behavior. We cannot presently differentiate between these alternatives but emphasize that the above possibilities are not mutually exclusive. Regardless, the parallelism between males and females suggests that some of the differences we observe are related to shifts in parental state that are independent of active behavioral caregiving.

While the overall pattern was the same, the absolute number of differentially expressed transcripts was greater in females. Furthermore, of the transcripts that differed between no care and tadpole transport versus egg care and tadpole transport, a smaller proportion was overlapping in males than females. In other words, the gene sets distinguishing care stages were more unique in males than females. Less overlap in differentially expressed transcripts between no care and tadpole transport versus egg care and tadpole transport in males may reflect the fact that, despite few significant expression differences between no care and egg care, these stages are more behaviorally distinct for males who are performing than females who are observing parental care. The relatively larger number of differentially expressed transcripts in females as compared to males is difficult to interpret but could arise from more extensive physiological differences across the reproductive cycle and/or differences associated with observing versus actively performing tadpole transport.

In addition to transcripts that differentiated males and females were those overlapping between sexes. Of the transcripts that were significantly differentially expressed during tadpole transport as compared to both no care and egg care, 45 were overlapping between males and females, all with expression changes in the same direction. We expect expression differences in the same direction if overlapping transcripts represents those involved in sex-independent transcriptional changes associated with tadpole transport (i.e. those with shared roles in parental care across sexes). One notable example is the increase in neuropeptide Y expression during tadpole transport in both caring males and observing female frogs. Neuropeptide Y (NPY) is most commonly associated with feeding behavior but several recent reviews have highlighted the adaptive trade-offs between feeding and social behavior and suggesting the circuitry mediating these behaviors is interconnected (Fischer and O’Connell, 2017; O’Rourke and Renn, 2015). Indeed, additional peptides and receptors commonly implicated in feeding behavior were also differentially expressed, including bombesin and cholecystokinin in males, and galanin and insulin-like growth factor receptors in females. Overall, these transcripts are promising candidates for future work examining sex-independent gene expression changes that re-tune neural circuits for parental care.

Finally, we note that we characterized whole-brain gene expression in parental males and their non-caregiving female partners across parental care stages. We chose a whole brain approach because the brain regions underlying parental behavior in amphibians were unknown at the time of this study, although these have recently been identified (Fischer et al. 2019). This approach likely masks more subtle, region-specific changes associated with parental behavior and does not allow us to distinguish transcript origin from neuronal versus glial cells; however, we emphasize that these limitations likely make our interpretations conservative.

### Neural activity patterns distinguish active and observing parents

We found that patterns of neural activity in transporting females were more similar to those we previously observed in transporting males than to those of non-caregiving female partners. Tadpole transporting males and females, but not non-caregiving females, had increased neural activity in the preoptic area, habenula, and suprachiasmatic nucleus. The preoptic area has been linked to parental behavior across vertebrates (reviewed in Fischer et al., 2019a; O’Connell and Hofmann, 2011), including across species and sexes in poison frogs (Fischer et al., 2019b). Our results here further support a role for the preoptic area in regulating tadpole transport behavior and suggest that tuning differences in preoptic area activity could mediate intra-specific plasticity as well as inter-specific variation in parental behavior between sexes. The lateral habenula is required for non-hormonally mediated maternal behavior in virgin female mice (Felton et al., 1998), providing a target for future work investigating hormone independent activation of tadpole transport. Given the tight link between sex typical hormone levels and social behavior, non-hormonally mediated mechanisms may be particularly interesting starting points for studies of sex-reversed behavior. Finally, the suprachiasmatic nucleus is most commonly associated with circadian rhythms (Kriegsfeld et al., 2002), and the increase in neural activity we observed here may be linked to timing dependent physiological and behavioral shifts across parental care stages. More experiments are needed to test these hypotheses and understand causal relationships among transcriptional, physiological, and neural activity changes.

In addition to the preoptic area, previous work identified the medial pallium (homolog of the mammalian hippocampus) as a core brain region associated with parental care in poison frogs (Fischer et al., 2019b). We observed increased neural activity in the medial pallium in tadpole-transporting males and females, as well as non-caregiving female partners. Notably, levels of neural activity were similarly elevated in non-caregiving females and non-tadpole transporting control females, but nonetheless lower than in actively transporting males or females. Given the important role of the medial pallium in spatial memory (Sotelo et al., 2016), we suggest that increased activity in non-caregiving and control *D. tinctorius* females is related to monitoring of male behavior. Although an additional increase in activity appears to be necessary for active performance of tadpole transport, this monitoring, in conjunction with hormone and brain gene expression changes, may prepare females to take over parental care. Additionally, in a study using the male uniparental poison frog *A. femoralis*, females did not cannibalize foreign (unrelated) egg clutches if they were swapped out with their own clutch, but did cannibalize their own and unrelated clutches if they were placed in a new location (Spring et al., 2019). This work in *A. femoralis* suggests that even when females do not directly perform parental duties, they have a precise spatial map of their egg clutch locations. These findings point to an important role for (spatial) memory in females independent of active parental behavior, lending support to our observation of increased medial pallial activity during tadpole transport in both active and observing *D. tinctorius*.

Overall, our findings in sex-reversed females suggest that hormonal changes across parental care stages are behavior-independent, while differences in neural activity are primarily linked to the active performance of parental care. We suggest that care-stage versus care-behavior specific associations between hormone levels, brain gene expression, and neural activity are linked to behavioral flexibility in parental care. As discussed above, hormonal changes in non-caregiving female partners may prime females to perform tadpole transport in the absence of their male mates. Although we did not measure brain gene expression in females performing sex-reversed transport, the similarities in expression patterns between males and females during sex typical care behavior suggests that transcriptional changes may serve a similar priming function. In contrast, patterns of neural activity associated only with the active performance of tadpole transport suggest that parental circuits present across sexes are primed by hormonal and gene expression changes, but that additional cues are necessary to activate these circuits and induce active parental care in females. This interpretation provides an explanation for parallel hormone and brain gene expression differences in caregiving males and their non-caregiving female partners, for an absence of hormonal differences between sex-reversed transporting and non-transporting control females, and for the potential of environmental conditions to induce sex-reversed parental care in this species.

### Conclusions

We found that similarities in neural activity were more closely related to active performance of parental behavior than hormone levels or whole brain gene expression, which were similar in parental males and their non-caregiving female partners. We suggest that hormonal and brain gene expression changes prime neural circuits for parental care, but additional factors trigger the changes in neural activity required for the ultimate performance of care behavior. Thus, the hormone and brain gene expression patterns we observe in non-caregiving *D. tinctorius* females may reflect their monitoring of male behavior and facilitate the changes in neural activity required when females take over parental duties to ensure the survival of their offspring. Continued exploration of behavioral flexibility in parental care will contribute to our understanding of how sex-specific behavioral patterns are coded in the brain, the conditions under which these patterns can be altered or reversed, and how circuits are evolutionarily tuned to give rise to species-specific care patterns.

## Acknowledgements

We thank the O’Connell Lab frog caretakers for help with animal care and Alexandre Roland for technical assistance. We also thank the members of the O’Connell Lab for discussions and Beau Alward and Kristina Smiley for comments on early versions of this manuscript.

## Funding

We gratefully acknowledge support from a Harvard University Bauer Fellowship (LAO), the International Society for Neuroethology Konishi Research Award (LAO), the Graduate Women in Science Adele Lewis Grant Fellowship (LAO), and an NSF postdoctoral fellowship (NSF-1608997 to EKF).

## Competing interests

The authors have no competing interests to declare.

## Notes

#### Summary of Updates

This manuscript has been revised per the comments of two anonymous reviewers.

## References

Adkins-Regan, E., 2005. Hormones and Animal Social Behavior. Princeton University Press, Princeton.

Albers, H.E., 2015. Species, sex and individual differences in the vasotocin/vasopressin system: relationship to neurochemical signaling in the social behavior neural network. Front. Neuroendocrinol. 36, 49–71.

Angelier, F., Chastel, O., 2009. Stress, prolactin and parental investment in birds: a review. Gen. Comp. Endocrinol. 163, 142–148.

Angelier, F., Wingfield, J.C., Tartu, S., Chastel, O., 2016. Does prolactin mediate parental and life-history decisions in response to environmental conditions in birds? A review. Horm. Behav. 77, 18–29.

Brown, R.E., 1985. Hormones and Paternal Behavior in Vertebrates. American Zoologist. 25, 895–910.

Caty, S.N., Alvarez-Buylla, A., Byrd, G.D., Vidoudez, C., Roland, A.B., Tapia, E.E., Budnik, B., Trauger, S.A., Coloma, L.A., O’Connell, L.A., 2019. Molecular physiology of chemical defenses in a poison frog. J. Expt. Biol. 222, jeb204149.

Clutton-Brock, T.H., 1991. The Evolution of Parental Care. Princeton University Press, Princeton.

Costantini, D., Marasco, V., Møller, A.P., 2011. A meta-analysis of glucocorticoids as modulators of oxidative stress in vertebrates. J. Comp. Physiol. B 181, 447–456.

da Silva Mota, M.T., Franci, C.R., de Sousa, M.B.C., 2006. Hormonal changes related to paternal and alloparental care in common marmosets (*Callithrix jacchus*). Horm. Behav. 49, 293–302.

Dumais, K.M., Veenema, A.H., 2016. Vasopressin and oxytocin receptor systems in the brain: Sex differences and sex-specific regulation of social behavior. Front. Neuroendocrinol. 40, 1–23.

Edelstein, R.S., Wardecker, B.M., Chopik, W.J., Moors, A.C., Shipman, E.L., Lin, N.J., 2015. Prenatal hormones in first-time expectant parents: Longitudinal changes and within-couple correlations. Am. J. Hum. Biol. 27, 317–325.

Felton, T.M., Linton, L., Rosenblatt, J.S., Morrell, J.I., 1998. Intact neurons of the lateral habenular nucleus are necessary for the nonhormonal, pup-mediated display of maternal behavior in sensitized virgin female rats. Behav. Neurosci. 112, 1458–1465.

Fischer, E.K., Nowicki, J.P., O’Connell, L.A., 2019a. Evolution of affiliation: patterns of convergence from genomes to behaviour. Philos. Trans. R. Soc. Lond. B. 374, 20180242.

Fischer, E.K., O’Connell, L.A., 2017. Modification of feeding circuits in the evolution of social behavior. J. Expt. Biol. 220, 92–102.

Fischer, E.K., Roland, A.B., Moskowitz, N.A., Tapia, E.E., Summers, K., Coloma, L.A., O’Connell, L.A., 2019b. The neural basis of tadpole transport in poison frogs. Proc. Roy. Soc. B. 286.

Fite, J.E., French, J.A., Patera, K.J., Hopkins, E.C., Rukstalis, M., Ross, C.N., 2005. Elevated urinary testosterone excretion and decreased maternal caregiving effort in marmosets when conception occurs during the period of infant dependence. Horm. Behav. 47, 39–48.

Fleming, A.S., Ruble, D., Krieger, H., Wong, P.Y., 1997. Hormonal and experiential correlates of maternal responsiveness during pregnancy and the puerperium in human mothers. Horm. Behav. 31, 145–158.

Fraser, B.A., Janowitz, I., Thairu, M., Travis, J., Hughes, K.A., 2014. Phenotypic and genomic plasticity of alternative male reproductive tactics in sailfin mollies. Proc. Biol. Sci. 281, 20132310.

Freedman, A.H., Clamp, M., Sackton, T.B., 2019. Error, noise and bias in de novo transcriptome assemblies. BioRxiv. https://doi.org/10.1101/585745

Gettler, L.T., McDade, T.W., Feranil, A.B., Kuzawa, C.W., 2011. Longitudinal evidence that fatherhood decreases testosterone in human males. Proc. Natl. Acad. Sci. 108, 16194–16199.

Harrison, F., Barta, Z., Cuthill, I., Székely, T., 2009. How is sexual conflict over parental care resolved? A meta-analysis. J. Evol. Biol. 22, 1800–1812.

Hashemian, F., Shafigh, F., Roohi, E., 2016. Regulatory role of prolactin in paternal behavior in male parents: A narrative review. J. Postgrad. Med. https://doi.org/10.4103/0022-3859.186389

Heatwole, H., Carroll, R.L., 2000. Amphibian biology. Surrey Beatty and Sons, Exeter.

Ketterson, E.D., Nolan, V., 1992. Hormones and life histories: an integrative approach. Am. Nat. 140 Suppl 1, S33–62.

Killius, A.M., Dugas, M.B., 2014. Tadpole transport by male *Oophaga pumlio* (Anura: Dendrobatidae): An observation and brief review. Herpetol. Notes 7, 747–749.

Knight, Z.A., Tan, K., Birsoy, K., Schmidt, S., Garrison, J.L., Wysocki, R.W., Emiliano, A., Ekstrand, M.I., Friedman, J.M., 2012. Molecular profiling of activated neurons by phosphorylated ribosome capture. Cell 151, 1126–1137.

Kohl, J., Autry, A.E., Dulac, C., 2017. The neurobiology of parenting: A neural circuit perspective. Bioessays 39, 1–11.

Kohl, J., Babayan, B.M., Rubinstein, N.D., Autry, A.E., Marin-Rodriguez, B., Kapoor, V., Miyamishi, K., Zweifel, L.S., Luo, L., Uchida, N., Dulac, C., 2018. Functional circuit architecture underlying parental behaviour. Nature 556, 326–331.

Kohl, J., Dulac, C., 2018. Neural control of parental behaviors. Curr. Opin. Neurobiol. 49, 116–122.

Kriegsfeld, L.J., LeSauter, J., Hamada, T., Pitts, S.M., Silver, R., 2002. Circadian Rhythms in the Endocrine System. Horm. Brain Behav. 2, 33–91.

Lavery, R.J., Reebs, S.G., 2010. Effect of Mate Removal on Current and Subsequent Parental Care in the Convict Cichlid (Pisces: Cichlidae). Ethology. 97, 65–77.

Love, M.I., Huber, W., Anders, S., 2014. Moderated estimation of fold change and dispersion for RNA-seq data with DESeq2. Genome Biol. 15, 550.

Magee, S.E., Neff, B.D., Knapp, R., 2006. Plasma levels of androgens and cortisol in relation to breeding behavior in parental male bluegill sunfish, *Lepomis macrochirus*. Horm. Behav. 49, 598–609.

McCormick, S.D., Bradshaw, D., 2006. Hormonal control of salt and water balance in vertebrates. Gen. Comp. Endocrinol. 147, 3–8.

Moore, F.L., Boyd, S.K., Kelley, D.B., 2005. Historical perspective: Hormonal regulation of behaviors in amphibians. Horm. Behav. 48, 373–383.

Norris, D.O., Lopez, K.H., 2010. Hormones and Reproduction of Vertebrates, Volume 2: Amphibians. Academic Press, Cambridge.

Numan, M., Insel, T.R., 2006. The Neurobiology of Parental Behavior. Springer Science & Business Media, Berlin.

O’Connell, L.A., Hofmann, H.A., 2011. The vertebrate mesolimbic reward system and social behavior network: a comparative synthesis. J. Comp. Neurol. 519, 3599–3639.

O’Connell, L.A., Matthews, B.J., Hofmann, H.A., 2012. Isotocin regulates paternal care in a monogamous cichlid fish. Hormones and Behavior. 61, 725–733.

Oliveira, R.F., Simões, J.M., Teles, M.C., Oliveira, C.R., Becker, J.D., Lopes, J.S., 2016. Assessment of fight outcome is needed to activate socially driven transcriptional changes in the zebrafish brain. Proc. Natl. Acad. Sci. 113, E654–61.

O’Rourke, C.F., Renn, S.C.P., 2015. Integrating adaptive trade-offs between parental care and feeding regulation. Curr. Opin. Behav. Sci. 6, 160–167.

Ouyang, J.Q., Sharp, P.J., Dawson, A., Quetting, M., Hau, M., 2011. Hormone levels predict individual differences in reproductive success in a passerine bird. Proc. Roy. Soc. B. 278, 2537–2545.

Pašukonis, A., Loretto, M.-C., Rojas, B., 2019. How far do tadpoles travel in the rainforest? Parent-assisted dispersal in poison frogs. Evolutionary Ecology. 33, 613–623.

Pereira, M., Ferreira, A., 2016. Neuroanatomical and neurochemical basis of parenting: Dynamic coordination of motivational, affective and cognitive processes. Horm. Behav. 77, 72–85.

Rilling, J.K., Mascaro, J.S., 2017. The neurobiology of fatherhood. Curr. Opin. Psychol. 15, 26–32.

Ringler, E., Pašukonis, A., Fitch, W.T., Huber, L., Hödl, W., Ringler, M., 2015. Flexible compensation of uniparental care: female poison frogs take over when males disappear. Behav. Ecol. 26, 1219–1225.

Rojas, B., 2014. Strange parental decisions: fathers of the dyeing poison frog deposit their tadpoles in pools occupied by large cannibals. Behav. Ecol. Sociobiol. 4, 551–559.

Rojas, B., Pašukonis, A., 2019. From habitat use to social behavior: natural history of a voiceless poison frog. PeerJ 7, e7648.

Roland, A.B., O’Connell, L.A., 2015. Poison frogs as a model system for studying the neurobiology of parental care. Curr. Opin. Behav. Sci. 6, 76–81.

Royle, N.J., Smiseth, P.T., Kölliker, M., 2012. The Evolution of Parental Care. Oxford University Press, Oxford.

Schindelin, J., Arganda-Carreras, I., Frise, E., Kaynig, V., Longair, M., Pietzsch, T., Preibisch, S., Rueden, C., Saalfeld, S., Schmid, B., Tinevez, J.-Y., White, D.J., Hartenstein, V., Eliceiri, K., Tomancak, P., Cardona, A., 2012. Fiji: an open-source platform for biological-image analysis. Nat. Methods 9, 676–682.

Smiseth, P.T., Dawson, C., Varley, E., Moore, A.J., 2005. How do caring parents respond to mate loss? Differential response by males and females. Animal Behaviour. 69, 551–559.

Sotelo, M.I., Daneri, M.F., Bingman, V.P., Muzio, R.N., 2016. Telencephalic Neuronal Activation Associated with Spatial Memory in the Terrestrial Toad Rhinella arenarum: Participation of the Medial Pallium during Navigation by Geometry. Brain Behav. Evol. 88, 149–160.

Spring, S., Lehner, M., Huber, L., Ringler, E., 2019. Oviposition and father presence reduce clutch cannibalism by female poison frogs. Front. Zool. 16, 8.

Storey, A.E., Walsh, C.J., Quinton, R.L., Wynne-Edwards, K.E., 2000. Hormonal correlates of paternal responsiveness in new and expectant fathers. Evol. Hum. Behav. 21, 79–95.

Summer, K., Weigt, L.A., Boag, P., Bermingham, E., 1999. The evolution of female parental care in poison frogs of the genus Dendrobates: Evidence from mitochondrial DNA sequences. Herpetologica 55, 254–270.

Townsend, D.S., Moger, W.H., 1987. Plasma androgen levels during male parental care in a tropical frog (Eleutherodactylus). Horm. Behav. 21, 93–99.

Wada, H., 2008. Glucocorticoids: mediators of vertebrate ontogenetic transitions. Gen. Comp. Endocrinol. 156, 441–453.

Wickham, H., 2009. ggplot2: Elegant Graphics for Data Analysis. Springer Science & Business Media.

Wilczynski, W., Lynch, K.S., O’Bryant, E.L., 2005. Current research in amphibians: Studies integrating endocrinology, behavior, and neurobiology. Horm. Behav. 48, 440–450.

Wingfield, J.C., Hegner, R.E., Dufty, A.M., Ball, G.F., 1990. The “Challenge Hypothesis”: Theoretical Implications for Patterns of Testosterone Secretion, Mating Systems, and Breeding Strategies. Am. Nat. 136, 829–846.

Wu, Z., Autry, A.E., Bergan, J.F., Watabe-Uchida, M., Dulac, C.G., 2014. Galanin neurons in the medial preoptic area govern parental behaviour. Nature 509, 325–330.

Wynne-Edwards, K.E., 2001. Hormonal Changes in Mammalian Fathers. Horm. Behav. 40, 139–145.

Young, L.J., Crews, D., 1995. Comparative neuroendocrinology of steroid receptor gene expression and regulation: Relationship to physiology and behavior. Trends Endocrinol. Metab. 6, 317–323.

Zahed, S.R., Kurian, A.V., Snowdon, C.T., 2009. Social dynamics and individual plasticity of infant care behavior in cooperatively breeding cotton-top tamarins. Am. J. Primatol. 72, 296–306.

